# Hierarchical spatial organization stabilizes algal-fungal living materials and enables sustained carbon fixation

**DOI:** 10.64898/2026.05.29.728589

**Authors:** Yujie Wang, Chi Hu, Jiaojiao Liu, Peng Chen, Xi Zeng, Zhaoxiang Yang, Ziyi Yu

## Abstract

Engineering stable multicellular living materials remains difficult because distinct partners require incompatible local microenvironments, while sustained cooperation depends on integration across multiple spatial scales. Here we show that hierarchical spatial organization stabilizes algal-fungal living materials and enables sustained carbon fixation. Compartmentalized seed-seedcase units create partner compatible local niches for *Chlorella vulgaris* and *Pleurotus ostreatus*, while fungal outgrowth bridges neighboring units into an integrated artificial lichen. This transition converts localized coexistence into a mechanically coherent assembly that sustains net CO_2_ drawdown together with O_2_ production in closed systems and restores carbon fixation activity after repeated CO_2_ replenishment. A printable formulation further enables macroscopic architecture with enhanced volumetric carbon-fixation performance. Transcriptomics reveals division of labor between algal carbon fixation and fungal redox and matrix support functions, whereas perturbation assays demonstrate functional robustness. These results establish hierarchical spatial organization as a design principle for stable cooperative living materials.

**Teaser:** Programmable algal-fungal assemblies turn spatial design into durable carbon capture.

## INTRODUCTION

Biological function often emerges from the spatial organization of distinct cellular partners into higher-order living systems *(1-6)*. Reconstituting this principle in engineered materials could enable living materials with emergent functions that isolated species cannot achieve alone*(7, 8)*. However, such systems require more than the simple co-presence of multiple organisms. Distinct partners must be organized across multiple spatial scales in ways that preserve compatible local microenvironments while still enabling exchange and integration at the level of the whole construct *(9-14)*. Therefore, a central challenge, is how to engineer multicellular living systems in which local compatibility and collective function can be sustained simultaneously.

Algal-fungal symbiosis provides a compelling model for engineering multicellular cooperation, with lichens representing its most successful natural example *(15-18)*. In these systems, photosynthetic and filamentous partners are functionally complementary, yet they depend on distinct and often incompatible local environments *(19, 20)*. Liquid co-culture offers little control over long-term spatial organization, whereas simple co-immobilization in solid or encapsulated formats can enforce proximity without reliably sustaining partner-specific niches, interfacial exchange, and construct-level integration *(21)*. As a result, although algal-fungal systems offer attractive opportunities for developing living materials with carbon-fixation capability, a general strategy for stabilizing such systems and sustaining function remains lacking *(22-24)*.

We therefore proposed that stabilizing such symbiosis would require hierarchical spatial control, such that local compartmentalization preserves partner-compatible niches and interfacial exchange, while connectivity between neighboring units enables construct-level integration *(25, 26)*. Extending this logic to macroscopic fabrication could further link symbiotic organization to manufacturable living materials. Herein, we show that hierarchical spatial organization stabilizes algal-fungal living materials and enables sustained carbon fixation. We construct compartmentalized seed-seedcase units that support *Chlorella vulgaris* (*C. vulgaris*) and *Pleurotus ostreatus* (*P. ostreatu*s) in distinct local niches, while fungal outgrowth bridges neighboring units into an integrated artificial lichen *(27, 28)*. Incorporation of gelatin further enables a printable formulation with programmable macroscopic architecture *(29)*. Together, these results identify hierarchical spatial organization as a strategy for converting algal-fungal coexistence into stable living materials with sustained carbon-fixation capability.

## RESULTS

### Local spatial compartmentalization establishes the basis for compatible algal-fungal living materials

Engineering algal-fungal living materials requires more than bringing the two partners into proximity, because stable coexistence depends on whether each organism can access a compatible local microenvironment while remaining coupled through interfacial exchange. We began to explore whether local spatial compartmentalization could provide compatible niches for *C. vulgaris* and *P. ostreatus* within a shared living unit. Sodium alginate (SA) was selected as the immobilization matrix because its mild ionic gelation provides hydrated porous networks with tunable transport and mechanical properties. Under these conditions, the two species responded differently to matrix composition: *C. vulgaris* formed larger aggregates in 0.5 wt% SA, whereas *P. ostreatus* grew more robustly in 2.5 wt% SA (Fig. S1). This different was consistent with the mechanical differences between the two formulations after gelation, with 2.5 wt% SA reaching a substantially higher modulus than 0.5 wt% SA. These contrasting responses were consistent with the marked mechanical difference between the two formulations, with 2.5 wt% SA exhibiting a substantially higher modulus than 0.5 wt% SA (Fig. S1), indicated that a uniform matrix would be unlikely to support balanced algal-fungal coexistence We therefore designed a compartmentalized “pomegranate” architecture in which microfluidically generated *C. vulgaris* hydrogel microparticles (*c*HMPs) served as the seed and a *P. ostreatus*-SA composite formed the surrounding seedcase (Fig. 1A). This seed-seedcase configuration spatially separated the two partners while preserving their material continuity within a shared scaffold. Because seed size is expected to affect both initial inoculum loading and transport accessibility, we compared seeds of 40, 200, and 1000 μm in diameter. *C. vulgaris* growth was strongly size dependent. Consistent with phenotypic observations, chlorophyll quantification showed that 200 μm seeds performed best, exceeding both 40 and 1000 μm seeds (Fig. S2). 200 μm seeds carried a substantially larger initial algal inoculum, increasing the likelihood of successful establishment. By contrast, although 1000 μm seeds further increased loading, their larger dimensions exacerbated light and mass-transfer gradients, leading to more heterogeneous growth between the periphery and the core. These results identified 200 μm as the optimal seed size for coordinating inoculum loading, transport accessibility, and growth uniformity within the compartmentalized unit (Fig. 1B).

**Fig. 1.**
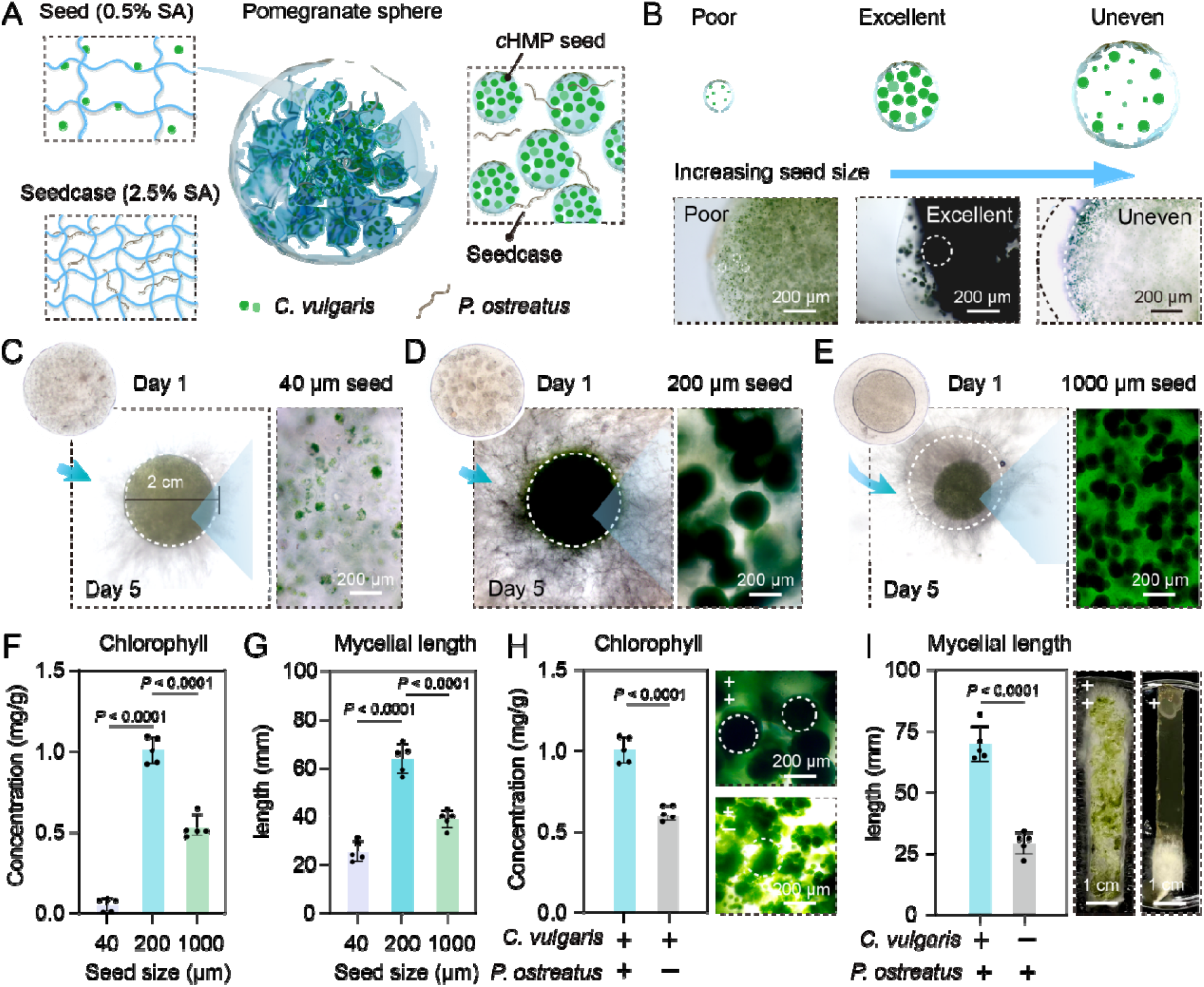
Local spatial compartmentalization establishes a compatible algal-fungal symbiotic unit. (**A**) Schematic illustration of the compartmentalized design for constructing algal-fungal pomegranate spheres, in which *C. vulgaris* was encapsulated as algal seeds in 0.5 wt% SA and *P. ostreatus* was incorporated into a fungal seedcase in 2.5 wt% SA. (**B**) Comparison of *C. vulgaris* growth in pomegranate spheres containing algal seeds of different sizes. (**C** to **E**)Representative images of pomegranate spheres containing algal seeds with diameters of 40 μm (**C**), 200 μm (**D**), and 1000 μm (**E**). *C. vulgaris* proliferated as dense aggregates within the algal seeds, while *P. ostreatus* mycelia penetrated the seedcase and extended outward. (**F**) Quantification of chlorophyll accumulation (**G**) Quantification of mycelial outgrowth. (**H**) Comparison of *C. vulgaris* growth between co-culture and monoculture groups. (**I**) Comparison of *P. ostreatus* mycelial outgrowth between co-culture and monoculture groups. Data in (**F**) to (**I**) are shown as mean ± SD (*n*=5 biological replicates). Statistical analysis was performed using one-way ANOVA. *P* values denote the statistical significance among the 40 μm seed, 1000 μm seed, and 200 μm seed groups. In (**F**) and (**G**), *P* values denote the statistical significance among the 40 μm seed, 1000 μm seed, and 200 μm seed groups. In (**H**), *P* values denote the statistical significance between the fungal group and the symbiotic group. In (**I**), *P* values denote the statistical significance between the algal group and the symbiotic group.

Using the optimized seed size, we assembled 200 μm algal seeds in 0.5 wt% SA with a fungal seedcase containing *P. ostreatus* in 2.5 wt% SA to form 2 mm pomegranate spheres. In this configuration, *C. vulgaris* proliferated as dense aggregates within the seed, whereas mycelia penetrated the seedcase and extended outward, generating a characteristic symbiotic interface in which a fraction of algal cells associated with the growing hyphae (Fig. S3). By day 5, pomegranate spheres with 200 μm seeds showed more than two-fold higher algal growth than those with 40 μm seeds or 1000 μm seeds (Fig. 1, C to E), together with markedly increased chlorophyll accumulation and mycelial outgrowth (Fig. 1, F and G). The co-incorporation of *C. vulgaris* and *P. ostreatus* promoted the growth of both partners. Consistent with this mutual growth enhancement (Fig.1, H and I), total protein accumulation also increased in the optimized formulation, consistent with greater overall biomass production (Fig. S4). Those results show that local spatial compartmentalization does not merely separate the two partners, but establishes a compatible and effective algal-fungal symbiotic unit in which algal proliferation, fungal extension, and overall material growth can proceed in a coordinated manner.

### Fungal bridging drives higher-order integration and sustained carbon fixation

A compartmentalized symbiotic unit provides the basis for local algal-fungal coexistence, but higher-order symbiosis also requires continuity across units. We next examined whether these units could be integrated into a higher-order living material. During co-culture, *P. ostreatus* progressively extended outward from individual pomegranate spheres and bridged neighboring particles, generating a continuous pomegranate artificial lichen (PomAL) rather than a loose aggregate of discrete units (Fig. 2A). This bridging process was accompanied by concurrent growth of the algal and fungal partners, indicating that interunit connectivity emerged together with continued symbiotic development.

**Fig. 2.**
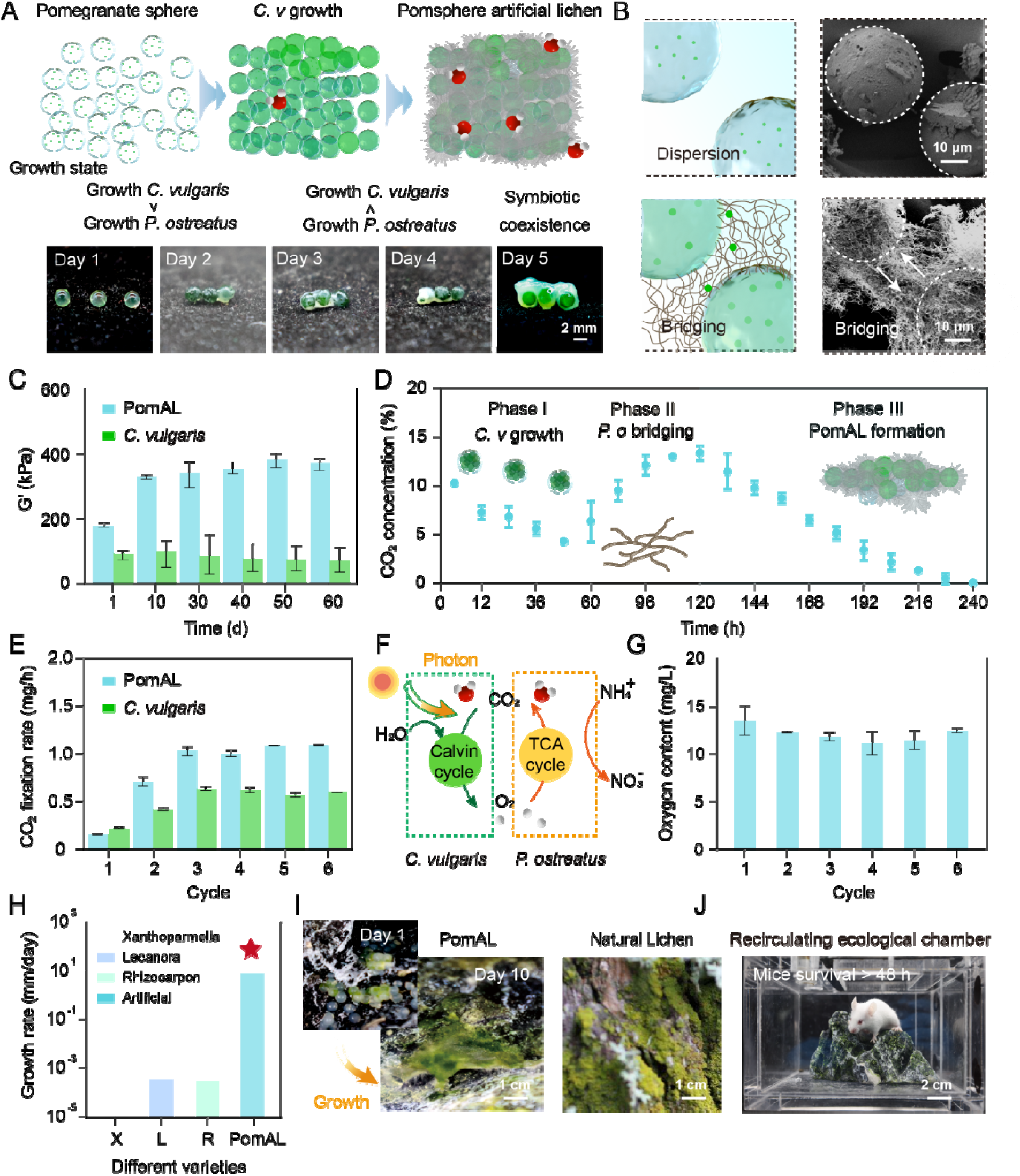
Fungal bridging drives the self-assembly of PomAL into an integrated living material for sustained carbon fixation. (**A**) Schematic illustration of intersphere bridging and self-assembly of PomAL. (**B**) SEM image of the bridged PomAL structure. (**C**) Time-dependent increase in the modulus of PomAL during growth. (**D**) Effect of PomAL on CO_2_ concentration in a closed environment. (**E**) Recoverable cyclic carbon-fixation performance of PomAL compared with *C. vulgaris*. (**F**) Schematic illustration of cyclic carbon fixation and oxygen production in the integrated assembly. (**G**) Repeated dissolved oxygen generation by PomAL. (**H**) Comparison of formation time among different lichen species. (**I**) Structural resemblance between PomAL and natural lichens. (**J**) Survival support provided by PomAL in a closed smoke-exposed chamber. Data in (**C**) to (**E**) and (**G**) are shown as mean ± SD (*n* = 3 biological replicates).

Morphological and mechanical analyses showed that this transition involved genuine structural integration rather than simple particle attachment. Microscopy and macroscopic imaging revealed that inter-sphere hyphae formed a continuous network across the assembly, whereas algal cells remained largely confined to the original compartmentalized domains and to interfacial regions associated with hyphal extension. PomAL therefore did not arise from collapse of the seed-seedcase architecture, but from hierarchical integration of preformed symbiotic units (Fig. 2, A and B, and Fig. S6). In this framework, compartmentalization and bridging played distinct but coordinated roles: the former preserved local spatial organization, whereas the latter established structural continuity across the construct. Mechanical measurements further supported this interpretation. As assembly progressed, the modulus of PomAL increased substantially relative to the initial sphere state, consistent with the development of a mechanically coherent mycelial network throughout the material, whereas the control lacking effective inter-sphere fungal growth showed little comparable strengthening (Fig. 2C). Fungal bridging was therefore not merely a morphological consequence of co-culture, but an organizational process that converted localized coexistence into an integrated multicellular living material.

We next explored whether this higher-order integration enabled functions beyond those of the individual components. When pomegranate spheres were placed in an airtight system, CO_2_ dynamics showed a staged transition during PomAL formation. During the first 48 h, rapid algal growth coincided with a decline in CO_2_ from about 10.2% to 4.2%. Between 48 and 96 h, accelerated mycelial growth and respiration produced a transient increase in CO_2_ to about 13.2%. After 96 h, once the PomAL structure had become established, the integrated assembly entered a sustained phase of net CO_2_ drawdown, with CO_2_ decreasing to near depletion by 128 h (Fig. 2D).

To determine whether the function was recoverable over repeated cycles, we repeatedly replenished CO_2_ and monitored the restoration of carbon-fixation activity. PomAL reproducibly restored net CO_2_ fixation after each recharge event and exhibited a substantially higher fixation rate than the algal-only control, reaching about 1.09 mg/h compared with 0.67 mg/h for *C. vulgaris* alone (Fig. 2E). Fungal bridging therefore transforms compartmentalized symbiotic units into a higher-order integrated living material that not only acquires structural continuity, but also supports sustained and recoverable carbon-fixation function. In parallel, the system continuously released O_2_ and maintained dissolved oxygen over multiple cycles (Fig. 2, F and G), indicating that the bridged material remained photosynthetically active while supporting fungal respiration. Notably, PomAL required only about one-thousandth of the time needed for natural lichens to establish stable structure and function (Fig. 2H), while exhibiting a broadly similar morphology (Fig. 2I), highlighting its rapid and biomimetic assembly advantage. Building on this, we further evaluated its environmental regulation potential in a closed system. A smoke-induced high-CO_2_ chamber was established, with mice used as physiological indicators to assess the effects of PomAL on air purification and survival support. PomAL effectively reduced chamber CO_2_ levels and improved the gaseous environment, thereby sustaining normal mouse survival (Fig. 2J, and Fig. S7).

### Architecture-level organization enhances volumetric carbon-fixation performance in printable living materials

Extension of the algal-fungal system from integrated assemblies to macroscopic constructs required a formulation that could preserve symbiotic organization while enabling programmable fabrication. To this end, we incorporated gelatin into the PomAL formulation to generate a printable artificial lichen (PAL) ink. The resulting formulation exhibited suitable rheological properties for extrusion-based fabrication and supported the concurrent growth of *C. vulgaris* and *P. ostreatus* on printed scaffolds (Fig. 3, A and B, and Fig. S8). This step enabled the algal-fungal system to be organized not only as compartmentalized and interconnected symbiotic units, but also as programmable macroscopic living materials. We then used PAL to construct porous scaffolds with different geometries and void sizes (Fig. 3C). These printed architectures provided a means to regulate light accessibility, mass transfer, and exposed surface area at the scale of the whole material. Among the tested designs, the small-pore structure (2 x 2 mm) structure supported the most favorable *C. vulgaris* and *P. ostreatus* growth and delivered the highest carbon-fixation performance (Fig. 3, C D, and F Fig. 3E Fig. S9), indicating that the geometry best balances transport accessibility with sufficient biomass retention. Denser structures likely restricted light penetration and diffusion, whereas more open structures reduced effective biomass density, making the small-pore configuration the optimal compromise between structural openness and functional loading.

**Fig. 3.**
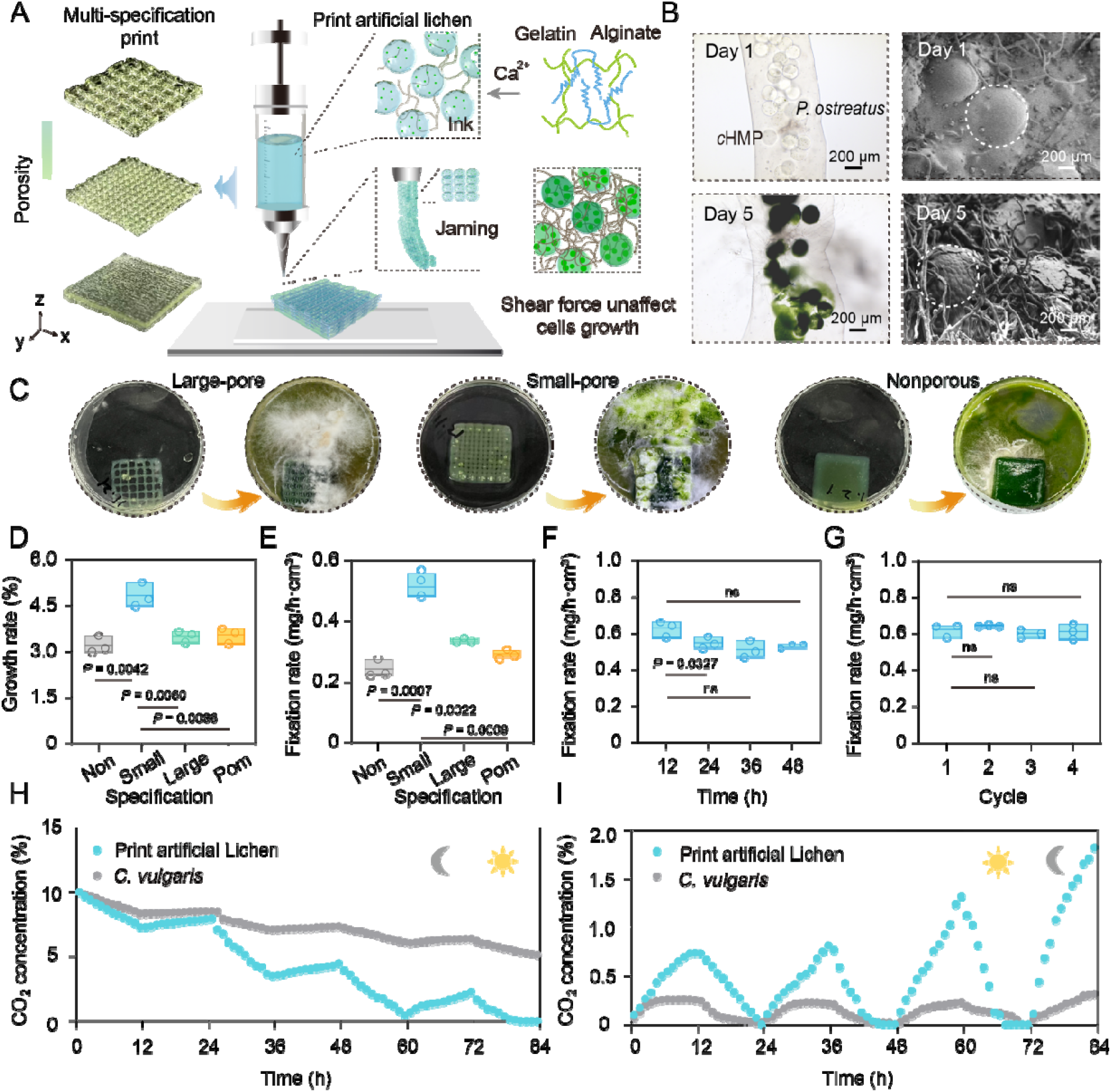
3D printing enables programmable fabrication of printed artificial lichen (PAL) and enhances its stable carbon-fixation performance. (**A**) Schematic illustration of PAL fabrication. (**B**) Microscopic and SEM characterization of PAL growth after fabrication. (**C**) Growth performance of 3D-printed PAL scaffolds with different pore sizes. (**D**) Comparison of the growth ratio of PAL scaffolds with different pore sizes. (**E**) Comparison of carbon-fixation rates of PAL with different pore sizes. (**F**) Stage-wise average carbon-fixation rate of small-pore PAL during a single fixation cycle. (**G**) Cyclic carbon-fixation rate of small-pore PAL over repeated fixation cycles. (**H**) Light-dark carbon-fixation performance of small-pore PAL under high-CO_2_ conditions. (**I**) Dark-light carbon-fixation performance of PAL under near-ambient CO_2_ conditions. Data in (**D**) and (**E**) are shown as mean ± SD (*n* = 3 biological replicates). Statistical analysis was performed using one-way ANOVA to compare each group with the small-pore group. *P* values denote statistical significance. Data in (**F**) and (**G**) are shown as mean ± SD (*n* = 3 biological replicates). Statistical analysis was performed using one-way ANOVA with each stage compared with stage I and each cycle compared with cycle 1.

To explore whether the architecture-level optimization also improved the stability of functional output, we monitored carbon fixation in the optimized small-pore PAL construct over time. During a 48 hour assay segmented into 4 h intervals, stage-wise fixation rates showed only minor variation, indicating stable short-term net CO_2_ fixation (Fig. 3F). Repeated CO_2_ replenishment further showed that PAL reproducibly maintained fixation activity over multiple cycles (Fig. 3G), demonstrating that the printed architecture supports repeatable functional performance rather than a transient single-cycle response.

To evaluate the adaptability of PAL under different carbon supplies and light regimes, we compared its carbon-fixation performance at elevated (10%) and near-ambient CO_2_ (0.04%) levels. Under high CO_2_, PAL sustained net carbon fixation during the light phase, whereas CO_2_ partially rebounded in the dark due to algal and fungal respiration (Fig. 3H). In the 12 h light/12 h dark regime, the dark phase likely provided both metabolic recovery and local CO_2_ accumulation from fungal respiration, contributing to a modest increase in fixation during the subsequent light period. Notably, PAL maintained the same overall trend of net CO_2_ uptake under near-ambient CO_2_, indicating stable function under low-carbon conditions (Fig. 3I). Overall, PAL consistently outperformed the algal-only system across CO_2_ conditions. Architecture-level organization is not merely a post-assembly processing step, but a third hierarchical layer of spatial organization that enhances volumetric carbon-fixation performance in printable algal-fungal living materials.

### Hierarchically organized living materials exhibit metabolic division of labor and functional robustness

Sustained carbon-fixation function in hierarchically organized algal-fungal living materials likely depends not only on structural organization, but also on coordinated metabolic specialization between the two partners. We therefore examined transcriptional reprogramming in PAL relative to algal and fungal monocultures. On the *C. vulgaris* side, enrichment in the Calvin cycle, pentose phosphate pathway, pyruvate metabolism, and glutathione metabolism (Fig. 4A), together with the upregulation of genes annotated as PRK, SBPase, CAB, FNR, GGDPR, and CPO, indicates enhanced light harvesting (Fig. 4B), reducing power supply, RuBP regeneration, and photosynthetic apparatus formation, thereby promoting CO_2_ fixation *(30-34)*. In parallel, the *P. ostreatus* side was enriched in glycolysis/gluconeogenesis, the pentose phosphate pathway, carbon metabolism, and amino acid metabolism (Fig. 4C), with upregulation of genes annotated as RKI1, FBA1, 6PGD, IDP1, UGD1, and GCV1, indicating strengthened carbon-flux reconfiguration, redox maintenance, and interfacial metabolic exchange (Fig. 4D). the upregulation of UGD1 likely promotes the production of extracellular polysaccharides and matrix components, creating a stable microenvironment that supports the attachment of microalgae. Through this interface, the microalgae can access some soluble metabolites, thereby enhancing their growth and CO_2_ fixation. At the same time, these changes also support fungal utilization of algal-derived photosynthetic products (such as sugars and sugar alcohols) and promote hyphal growth and bridging *via* TCA cycle activity. Active fungal nitrogen metabolism and IAA-related pathways may further improve the nutrient status and growth of the microalgae. Together, these interactions establish a positive-feedback symbiosis in PAL, in which algal carbon supply, fungal metabolic integration, and reciprocal support jointly enhance overall carbon-fixation performance (Fig. 4E).

**Fig. 4.**
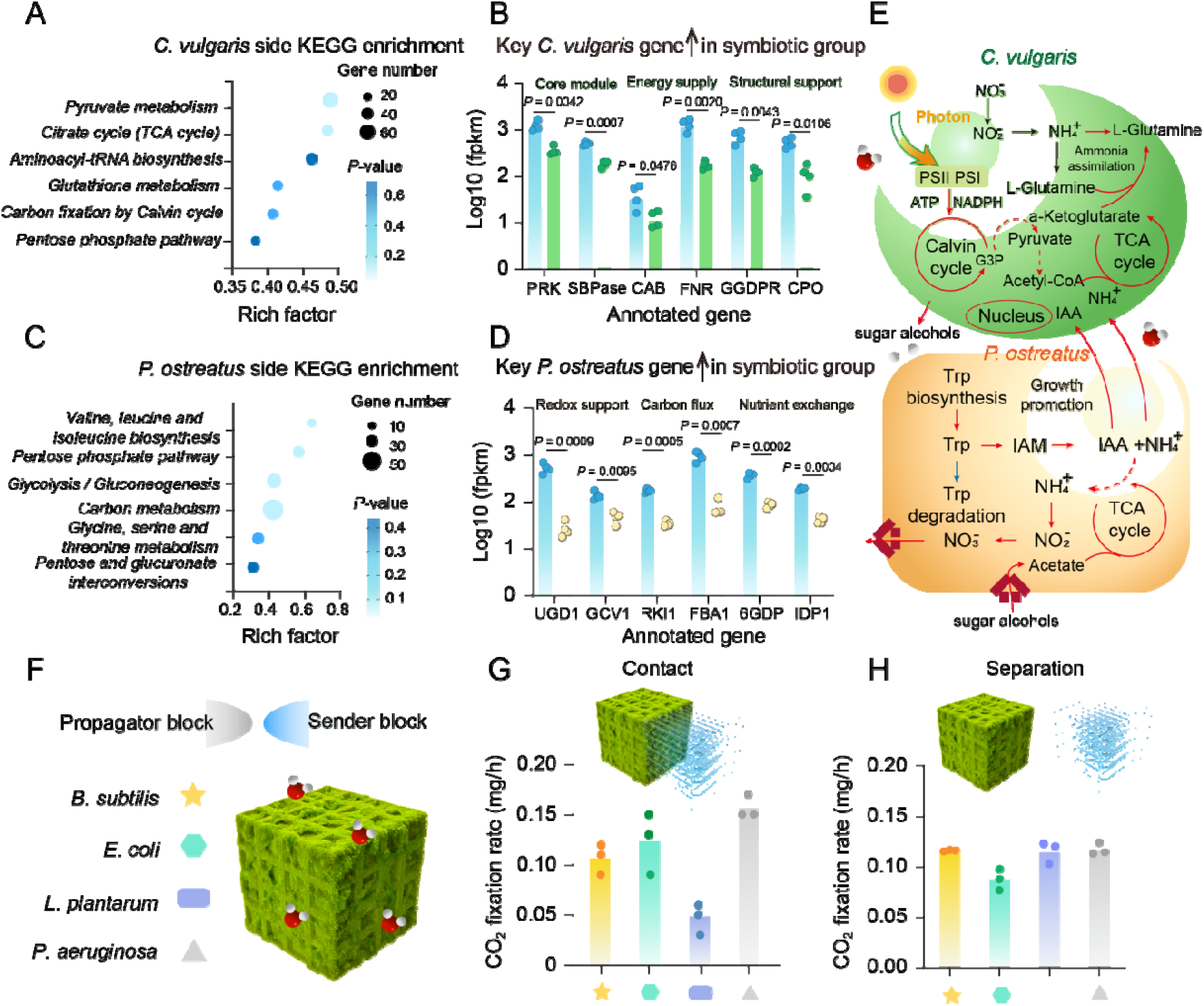
Molecular basis of symbiosis-enhanced carbon fixation and microbial perturbation resistance in PAL. (**A**) KEGG enrichment analysis of *C. vulgaris* side in the symbiotic group.(**B**) Upregulated genes on the *C. vulgaris* side in the symbiotic group. PRK (phosphoribulokinase), SBPase (sedoheptulose-1,7-bisphosphatase), CAB (chlorophyll a/b-binding protein), FNR (ferredoxin-NADP+ reductase), GGDPR (geranylgeranyl diphosphate reductase), CPO (coproporphyrinogen III oxidase). (**C**) KEGG enrichment analysis of the *P. ostreatus* side in the symbiotic group. (**D**) Upregulated genes on the *P. ostreatus* side in the symbiotic group. UGD1 (UDP-glucose dehydrogenase), GCV1 (aminomethyltransferase / glycine cleavage system T protein), RKI1 (ribose-5-phosphate isomerase), FBA1 (fructose-1,6-bisphosphate aldolase), 6PGD (6-phosphogluconate dehydrogenase) IDP1 (mitochondrial NADP-dependent isocitrate dehydrogenase). (**E**) Schematic illustration of the symbiotic mechanism underlying enhanced carbon fixation. (**F**) Schematic illustration of PAL resistance to microbial perturbation. (**G**) Carbon-fixation performance of PAL under direct-contact microbial perturbation. (**H**) Carbon-fixation performance of PAL under indirect microbial perturbation. Data in (**B**) and (**D**) are shown as mean ± SD (*n* = 4 biological replicates). Data in (**G**) and (**H**) are shown as mean ± SD (*n* = 3 biological replicates). Statistical analysis was performed *via* one-way ANOVA. *P* values denote the statistical significance of comparisons between the algal group and the algal component of the symbiotic group, and between the fungal group and the fungal component of the symbiotic group.

We further examined whether this hierarchically organized system could maintain function under external microbial challenge (Fig. 4F). To reconstruct a controllable multi-microbial cohabitation interface, PAL was used as a sender block and placed in direct or indirect contact with propagator blocks containing *Bacillus subtilis* (*B. subtilis*) *Escherichia coli* (*E. coli*), *Lactobacillus plantarum* (*L. plantarum*), or *Pseudomonas aeruginosa* (*P. aeruginosa*). Under direct contact, all exogenous microbes reduced carbon-fixation activity, with *E. coli* exerting the strongest inhibition, lowering the fixation rate to about 0.087 mg/h (Fig. 4G, and Fig. S10). Under indirect contact, the response became more differentiated: PAL was most sensitive to *L. plantarum*, whereas exposure to *B. subtilis, E. coli*, or *P. aeruginosa* caused only moderate reductions and still allowed net carbon fixation to be maintained (Fig. 4H, and Fig. S11). These distinct response profiles suggest that direct perturbation is dominated by interfacial attachment and near-field competition, whereas indirect perturbation is shaped more strongly by diffusible changes in the shared microenvironment, including acidification and redox imbalance.

Hierarchically organized algal-fungal living materials therefore exhibit both metabolic specialization and functional robustness. Spatial organization not only partitions structural roles across multiple scales, but also supports a coordinated division of labor that sustains carbon input, metabolic conversion, and material growth. At the same time, the PAL architecture buffers microbial perturbation sufficiently to preserve core function across a range of exogenous interactions. This combination of specialization and robustness helps explain why sustained carbon-fixation activity can be maintained in the engineered system.

## DISCUSSION

In summary, we have developed a hierarchically organized algal-fungal living material that stabilizes multicellular cooperation and sustains carbon-fixation function through coordinated spatial organization of symbiotic partners. Through compartmentalized seed-seedcase units, fungal bridging between neighboring particles, and architecture-level fabrication, this system provides partner-compatible niches, supports construct-level integration, and enables sustained net CO_2_ drawdown together with O_2_ production and enhanced volumetric carbon-fixation performance in printable constructs. Transcriptomic and perturbation analyses further show that this spatial organization is coupled to metabolic division of labor and functional robustness. More broadly, our results identify hierarchical spatial organization as a practical design principle for engineering cooperative living materials from partners with incompatible local requirements.

## MATERIALS AND METHODS

### Materials

Sodium alginate (MW, 222 kDa, Shanghai Macklin), calcium chloride (Aladdin), tryptone (Shanghai Macklin), yeast extract (Aladdin), sodium chloride (Shanghai Macklin), peptone (Aladdin), beef extract (Shanghai Macklin),(Shanghai Macklin), yeast extract (Aladdin), glucose (Aladdin), dipotassium hydrogen phosphate (K_2_HPO_4_; Shanghai Hushi), diammonium hydrogen citrate (Aladdin), sodium acetate (Shanghai Macklin), magnesium sulfate (Sigma), manganese sulfate (Sigma), Tween 80 (Shanghai Macklin), fluorescein isothiocyanate (FITC) (Shanghai Macklin), kanamycin (Shanghai Macklin).

### Cell preparation

#### Biological strains

*C. vulgaris* (Shanghai Guangyu Biotechnology Co., Ltd.; Cat. No. *C. vulgaris* GY-D19). *P. ostreatus* (Jingbao Bio). *B. subtilis* (gift by the Jiang Min group, Nanjing Tech University). *E. coli* ATCC 25922-sfGFP. *L. plantarum* (Culture Collection of Jiangnan University; strain *L. plantarum* CCFM8610, ST-III). *P. aeruginosa* (Beina Bio-Henan Engineering Technology Research Center of Industrial Microbial Strains; Cat. No. BNCC337940).

### Microbial culture conditions

*P. ostreatus*: *P. ostreatus* was cultured on PDA agar. The medium was autoclaved at 115°C for 20 min. A 2 x 2 cm mycelial plug was excised with a sterile scalpel and transferred onto fresh PDA plates. Plates were incubated unsealed at 25-28°C for 4 weeks. *C. vulgaris*: *C. vulgaris* was maintained on BG11 medium. BG11 was autoclaved at 121°C for 15 min, cooled to 40°C, and supplemented with kanamycin (working concentration: 10 ug/mL). After mixing, the medium was poured into sterile Petri dishes. Algal cells from liquid culture were streaked onto BG11 plates using a sterile inoculating loop and incubated in a constant-temperature chamber at 25°C under continuous illumination (6000 lux) at 65% relative humidity for 25 days. Single colonies were then picked and inoculated into BG11 liquid medium (100 mL H_2_O + 0.17 g BG11), followed by cultivation at 25°C under continuous illumination (6000 lux) with shaking at 150 rpm for 7 days. *B. subtilis*: A 100 μL aliquot of glycerol stock was inoculated into 5 mL LB broth and cultured at 37°C, 200 rpm for 24 h. *E. coli*: A 100 μL aliquot of glycerol stock was inoculated into 5 mL LB broth and cultured at 37°C, 200 rpm for 24 h. *L. plantarum*: A 100 μL aliquot of glycerol stock was inoculated into 5 mL MRS broth and cultured at 37°C, 200 rpm for 24 h. *P. aeruginosa*: A 100 μL aliquot of glycerol stock was inoculated into 5 mL LB broth and cultured at 37°C, 200 rpm for 24 h.

### Microfluidic fabrication of alginate microgels

To preserve microalgal viability, an ion-exchange strategy was used to prepare SA microgels instead of acetic-acid-induced crosslinking. A highly monodisperse alginate microbead emulsion was generated in a microfluidic device using three phases: water phase 1 consisting of an 80 mM Ca-EDTA chelate solution mixed with SA; water phase 2 consisting of an 80 mM Zn-EDDA chelate solution mixed with SA and an oil phase containing 2% (v/v) Drop-Surf in HFE-7500. Algal paste was added to each water phase to reach a final OD_680_ of 1.5. The flow rates of water phase 1, water phase 2, and the oil phase in the microfluidic device were set to 5, 5, and 20 μL/min, respectively. Microbeads were collected into centrifuge tubes and allowed to gel for 10 min. The beads were then washed sequentially with demulsifier and culture medium to remove the fluorinated oil. Microfluidic chips with different channel dimensions were used to generate *c*HMPs of defined diameters (e.g., 40 μm).

### Preparation of pomegranate spheres

Pomegranate spheres were formed *via* ionic crosslinking between SA and CaCl_2_. The setup consisted of a vertically mounted syringe pump and a Petri dish containing CaCl_2_ solution. Briefly, 0.1 g of *P. ostreatus* mycelium was homogenized with a sterile grinding rod and mixed thoroughly with 4 mL of 2.5% (w/v) SA, serving as the “rind” component. The collected *c*HMPs were then mixed with the rind matrix at 1:1 (v/v) to obtain the pomegranate-sphere ink. The ink was loaded into a 1 mL syringe fitted with a 21 G luer-lock dispensing needle (0.56 mm inner diameter, 0.82 mm outer diameter). The syringe pump was set to an extrusion rate of 10 μL/min, and droplets were dispensed into 4% (w/v) CaCl_2_ solution and crosslinked for 5 min. The resulting pomegranate spheres were cultured at 25°C in a mixed liquid medium (BG11:PDA = 5:1, v/v). With increasing culture time, the spheres transitioned from light green to dark green due to algal proliferation. Mycelia traversed the interior and extended outward, intertwining to bridge neighboring spheres and ultimately forming an integrated construct.

### Growth characterization of pomegranate spheres

#### Growth ratio

For 200 μm seed pomegranate spheres, excess moisture was removed with sterile filter paper prior to weighing. The growth ratio was calculated as follows:

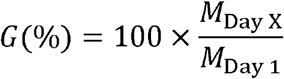

G(%): growth rate; M_Day x_: the weight measured on day x (x > 1) Under identical experimental conditions, 40 μm seed pomegranate spheres were processed and measured in parallel as controls for growth comparison.

#### Mycelial length measurement

A single 200 μm seed pomegranate sphere was placed at one edge of a PDA agar plate. Mycelia emerged from the sphere and extended onto the PDA surface for further growth, and the outgrowth length was measured. 40 μm seed pomegranate spheres were tested in parallel to compare mycelial outgrowth length. Protein content measurement (BSA assay). 200 μm seed pomegranate spheres were resuspended in 2 mL 1xPBS and washed three times. The samples were homogenized at 18,000 rpm for 10 min and subjected to three freeze-thaw cycles in liquid nitrogen to ensure complete cell disruption. After centrifugation, the supernatant was collected and absorbance was measured at 595 nm using a microplate reader. A bovine serum albumin (BSA) standard curve was used for quantification, with an equal volume of 1xPBS as the blank control. 40 μm seed pomegranate spheres were processed and measured in parallel under identical conditions to compare protein content.

#### Chlorophyll content measurements

Approximately 60 mg of 200 μm seed pomegranate spheres was resuspended in 0.5 mL methanol and incubated in a 60°C water bath for 30 min. The samples were then centrifuged at 3,000 rpm for 5 min. The supernatant was collected and absorbance was measured at 665, 652, and 470 nm using a microplate reader. Total chlorophyll and carotenoid concentrations were calculated according to the following equations:

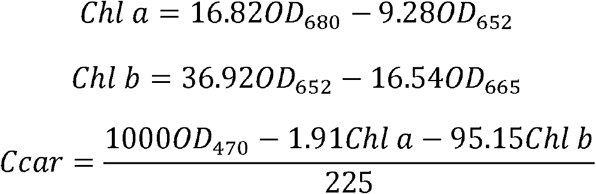

Where Chl a and Chl b denote chlorophyll a and chlorophyll b contents, respectively, and Ccar denotes carotenoid content. Under identical experimental conditions, pomegranate sphere samples with 40 μm seeds were processed in parallel to determine chlorophyll and carotenoid contents as controls.

### SEM characterization of pomegranate spheres

Samples were prepared by graded dehydration. 200 μm seed pomegrante spheres were resuspended in 2 mL 1xPBS and washed three times, then fixed in 2.5% (w/v) glutaraldehyde at 4°C overnight, followed by rinsing with 0.1 M PBS buffer. Dehydration was carried out by sequential immersion in graded ethanol solutions (50%, 70%, 80%, 90%, 95%, and 100%). Samples were freeze-dried for 12 h, mounted on SEM stubs, and sputter-coated with gold (8 mA, 60 s). Morphology was examined using a scanning electron microscope (Sigma 360, Carl Zeiss).

### Stress-strain measurements

200 μm seed pomegrante spheres collected at different culture stages were tested on a rheometer (Thermo Electron D-76227) equipped with a P8/Ti rotor (8 mm). Oscillatory strain-amplitude sweeps were performed over a strain range of γ = 0.0001-10 at 25°C. The control group consisted of pomegranate spheres containing only microalgae.

### CO_2_ fixation measurements of pomegranate spheres

Ten grams of freshly prepared pomegranate spheres were loaded into a test vessel (150 mL vacuum filtration flask; bottom diameter 7 cm, height 12 cm, neck height 3.5 cm, mouth diameter 1.55 cm; total volume ≈ 0.21 L). A CO_2_ probe (CARBOCAP-FMP251) was inserted through the neck and positioned 5-7 cm away from the sample. The flask was sealed, flushed with CO_2_ for 10 min, and timing was initiated once the CO_2_ reading stabilized. For cyclic CO_2_ fixation tests, the sample was kept in the sealed vessel and repeatedly charged with the same volume of CO_2_, while monitoring CO_2_ concentration changes over time in the closed environment.

Calculation of CO_2_ fixation amount:

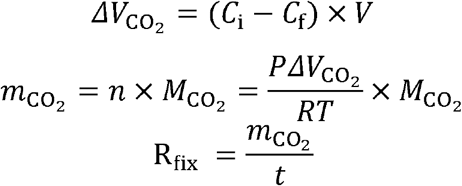

C_i_ : initial CO_2_ concentration; C_f_ : final CO_2_ concentration.

### Dissolved oxygen measurements of pomegranate spheres

Ten grams of freshly prepared pomegranate spheres were placed in a test vessel (cylindrical bottle; 4 cm diameter, 10 cm height) and supplemented with 20 mL of mixed culture medium. An oxygen probe was immersed in the medium, and the vessel was sealed with a parafilm membrane. A gas inlet needle was extended into the liquid and NL was bubbled to remove dissolved O_2_. Monitoring was initiated once the displayed dissolved oxygen reached 0%.

### PomAL mountain experiment

Rocks were pre-soaked in mixed culture medium and then removed. 5 g of freshly prepared pomegranate spheres were randomly dispersed over the rock surface and incubated in a sealed sterile environment at 25°C under continuous illumination (24 h) and 60% relative humidity. During cultivation, mixed liquid medium was sprayed every 2 days onto areas containing pomegranate spheres. PomAL formed after 10 days.

### Closed-room experiment

A closed room (140 × 100 × 150 mm) was fabricated from acrylic and equipped with four ports: (1) smoke inlet port (4.6 mm diameter), (2) exhaust port (4.6 mm diameter), (3) CO_2_ detection port (4.6 mm diameter), and (4) animal entrance (80 mm x 65 mm). The experimental group included the closed room, the PomAL Mountain, and one 4-5-month-old BALB/c mouse; the control group included the closed room, a blank mountain construct and one 4-5-weeks-old BALB/c mice. A cigarette (Yuxi brand, Yunnan Hongta Group Yuxi Cigarette Factory) was ignited, and the burning end was inserted into port 1. When the CO_2_ concentration in the chamber reached 5%, the cigarette was extinguished and port 1 was sealed. Mouse activity in the closed room was then observed. Food and jelly were provided in both the experimental and control groups to meet basic survival needs. Male BALB/c mice (18-22 g, 4-5 weeks old) were purchased from Jiangsu Huachuang Xinnuo Pharmaceutical Technology Co., Ltd. (Taizhou, China). All mice were housed under specific pathogen-free (SPF) conditions with strictly controlled environmental parameters (22 ± 2°C, 50 ± 5% relative humidity) on a 12-hour light/dark cycle. Animals had ad libitum access to standard chow and water and were housed in identical cages with wood shavings as bedding. All animal experiments were approved by the Institutional Animal Care and Use Committee of China Pharmaceutical University (approval no. YSL-202604050) and complied with ARRIVE guidelines. Throughout the study, efforts were made to minimize animal use and alleviate potential discomfort.

### Construction and performance evaluation of PAL

The pomegranate-sphere ink was supplemented with 2% (w/v) type A gelatin to obtain the printing formulation for artificial lichen, termed PAL ink. Rheological characterization of PAL ink. The rheological properties of PAL ink were measured using a stress-controlled rheometer (HAAKE Rheostress 1, Thermo Scientific, USA) equipped with a 35 mm conical plate (gap: 100 μm) at 25°C. Flow curves were obtained in shear rate-controlled mode over a shear-rate range of 0.02-100 s^-1^. Dynamic oscillatory strain-amplitude sweeps were performed at 1 Hz, and frequency sweeps were conducted over 0.1-100 rad/s.

Extrusion-based printing of PAL ink. PAL ink was printed using an extrusion bioprinter (EFL-BP-6600) equipped with a low-temperature printhead and a cooled printing stage. Structures were fabricated layer-by-layer with a layer height of 0.2 mm. The printing speed was set to 1000 mm/min with a 90° deposition angle, and the perimeter (edge) speed was 1500 mm/min. The printed scaffolds were immersed in 4% (w/v) CaCl_2_ solution for 5 min to crosslink. All printing geometries were designed in Autodesk Fusion 360. Porous-architecture printing. One milliliter of PAL ink was used to print square scaffolds (2.4 x 2.4 cm) with different void sizes. For the nonporous scaffold, the void size was 0 x 0 mm with ∼12 layers printed. For the small-pore design, the void size was 2 x 2 mm with ∼28 layers printed. For the large-pore design, the void size was 4 x 4 mm with ∼48 layers printed. CO_2_ fixation measurements of PAL. A mixed solid medium was placed at the bottom of the test vessel, and the printed scaffold was positioned inside. The vessel was then sealed, CO_2_ was introduced through a gas inlet needle, and changes in the CO_2_ concentration inside the vessel were monitored.

### PAL transcriptomic sequencing

#### Sample preparation

The experimental group consisted of 2.5 g the symbiotic group beads. Control 1 consisted of 2.5 g algal beads (0.5% (w/v) SA, OD_680_ of 1.5), prepared by directly dripping the algal-alginate precursor into 4% (w/v) CaCl_2_ to form gel beads (2 mm diameter). Control 2 consisted of 2.5 g *P. ostreatus* beads, prepared by homogenizing 0.1 g *P. ostreatus* mycelium with 4 mL of 2.5% (w/v) SA and directly dripping the mixture into 4% (w/v) CaCl_2_ to form gel beads (2 mm diameter). All groups were cultured in mixed medium for 10 days at 25°C under continuous illumination. Four biological replicates were prepared for each group. After culture, samples were resuspended in 3 mL 1xPBS and washed 5 times, then resuspended in 2 mL TRIzol (Yeasen). Total RNA was extracted using RNA-easy Isolation Reagent (Vazyme) and quantified using the Qubit RNA HS Assay Kit on a Qubit 2.0 fluorometer (Life Technologies, CA, USA). Transcriptome sequencing was carried out on an Illumina NovaSeq 6000 platform (Illumina, San Diego, CA, USA) at Metware Biotechnology Co., Ltd. (Wuhan, China).

### Interactions between PAL and multiple microbes

#### Construction of microbial scaffolds

The four bacterial cultures were adjusted to OD_500_ of 0.5 in liquid medium, centrifuged at 3,000 rpm for 3 min to remove the supernatant, and resuspended in 2.5% (w/v) SA. Type A gelatin (2% (w/v)) was then added to obtain the microbial ink. The microbial ink was used to print cubic scaffolds with 2 x 2 mm voids and 28 layers, serving as the Sender block. Cubic scaffolds (2 x 2 mm voids) printed from PAL ink served as the Propagator block.

#### Spatial sensing configurations

For separated sensing, the Sender and Propagator blocks were placed on partitioned media within the test vessel, with a 4 cm gap between blocks. For co-localized sensing, the Sender and Propagator blocks were placed on the same medium region, with their side surfaces in direct contact.

#### Partitioned media preparation

Because the media requirements of PAL and the bacteria were incompatible, partitioned media were used. The PAL region contained mixed solid medium, whereas the microbial region contained the corresponding solid medium for each bacterium (LB for *B. subtilis*, LB for *E. coli*, MRS for *L. plantarum*, and LB for *P. aeruginosa*).

#### Co-localized medium

For the co-localized medium, mixed medium and the corresponding bacterial medium were combined at 1:1 (v/v) with 2 wt% agar.

#### CO_2_ fixation measurements under coexistence with multiple microbes

Partitioned media were set in the test vessel, and PAL and the microbial scaffolds were placed in their respective regions. The probe was connected, CO_2_ was introduced for 10 min, the vessel was sealed, and the CO_2_ concentration was then monitored.

### Statistical analysis

Statistical analyses were performed using GraphPad Prism 9.0 (GraphPad Software). Data are presented as mean ± SD. Sample size (*n*), statistical methods, and exact *P* values are indicated in the corresponding figure legends or main text where appropriate. For comparisons among multiple groups, one-way analysis of variance (ANOVA) was used unless otherwise specified. For comparisons between two groups, unpaired two-tailed Student’s t test was used unless otherwise indicated. A value of *P* < 0.05 was considered statistically significant. Unless otherwise stated, quantitative experiments were performed with at least three biological replicates, and qualitative experiments were independently repeated at least three times.

## Acknowledgments

This work was supported the National Key Research and Development Program of China (2024YFA0919100) and National Natural Science Foundation of China (T2322011, 22278214, 52003119).

## Author contributions

Conceptualization: C. H., Z. Y. Methodology: Y. W., C. H., J. L., P. C.

Investigation: Y. W., J. L.

Visualization: Z. Y., C. H., Y. W.

Supervision: C. H., Z. Y.

Writing-original draft: Z. Y., C. H., Y. W.

Writing-review & editing: Z. Y., C. H., X. Z., Z. Y.

